# Fast and Flexible GPU Accelerated Binding Free Energy Calculations within the AMBER Molecular Dynamics Package

**DOI:** 10.1101/247692

**Authors:** Daniel J. Mermelstein, Lin Charles, Nelson Gard, Kretsch Rachael, J. Andrew McCammon, Ross C. Walker

## Abstract

Alchemical free energy calculations (AFE) based on molecular dynamics (MD) simulations are key tools in both improving our understanding of a wide variety of biological processes and accelerating the design and optimization of therapeutics for numerous diseases. Computing power and theory have, however, long been insufficient to enable AFE calculations to be routinely applied in early stage drug discovery. One of the major difficulties in performing AFE calculations is the length of time required for calculations to converge to an ensemble average. CPU implementations of MD based free energy algorithms can effectively only reach tens of nanoseconds per day for systems on the order of 50,000 atoms, even running on massively parallel supercomputers. Therefore, converged free energy calculations on large numbers of potential lead compounds are often untenable, preventing researchers from gaining crucial insight into molecular recognition, potential druggability, and other crucial areas of interest. Graphics Processing Units (GPUs) can help address this. We present here a seamless GPU implementation, within the PMEMD module of the AMBER molecular dynamics package, of thermodynamic integration (TI) capable of reaching speeds of >140 ns/day for a 44,907-atom system, with accuracy equivalent to the existing CPU implementation in AMBER. The implementation described here is currently part of the AMBER 18 beta code and will be an integral part of the upcoming version 18 release of AMBER.

## Introduction

Computational chemists have for some time been seeking to correctly and universally predict the answer to the question “How well will ligand X bind to protein Y?” Couched in this is the implicit assumption that our predictive tool will be able to answer this question in a reasonable amount of time. For years, computational chemists have been forced to strike a difficult balance between speed and accuracy, often being forced to rely on faster, less accurate methods because (TI) and related Free Energy Perturbation (FEP) methods were simply too computationally intensive. However, the ceiling on the applicability of docking and MM/PBSA for use in making quantitative predictions of ligand binding, possibly the most relevant aspect of free energy in the pharmaceutical industry, has quickly become apparent.^1^ This has led to a renewed interest in improving slower but potentially more accurate methods such as TI. One of the primary difficulties which TI suffers from is a severe sampling limitation.^2^ This limitation is the result of several factors including the difficulty in sampling the relevant states,^3^ potentially dozens of unphysical intermediate states that must be simulated even for systems as small as host-guest systems,^4^ and the complexity and cost of the underlying molecular dynamics based algorithm. Thus, either hundreds of CPU cores and/or weeks of simulation time are required to obtain results, within acceptable error limits, for a single ligand-protein system in a drug design setting. This is untenable in an industrial setting where high-throughput is integral to success. The purpose of this work is to help address the throughput and finite sampling problem in TI using cost effective consumer hardware.

While there have been many different approaches to addressing the issue of sufficient sampling in TI calculations^5–7^ the most straightforward manner of improving on the sampling is to write faster code. This paper details a substantially faster implementation of alchemical transformations using cost effective, NVIDIA GeForce graphics processors (GPUs). This code is fully implemented within the AMBER molecular dynamics package,^8–10^ and takes direct advantage of the existing, highly efficient, GPU support.^11–13^ The implementation described here will be released as an integral part of the GPU accelerated PMEMD program in the upcoming AMBER 18 software release scheduled for Q2 2018. It is currently part of the AMBER 18 Beta code and a patch against AMBER 17 is available by request from the corresponding author.

## Theory and Methods

### Model Calculations

The goal of this paper is to demonstrate the efficacy and numerical precision of our new GPU TI implementation in AMBER. We have chosen two types of calculations for this purpose (Figure 1). The first, a free energy of solvation, the second, a binding affinity calculation.

**Figure 1:**
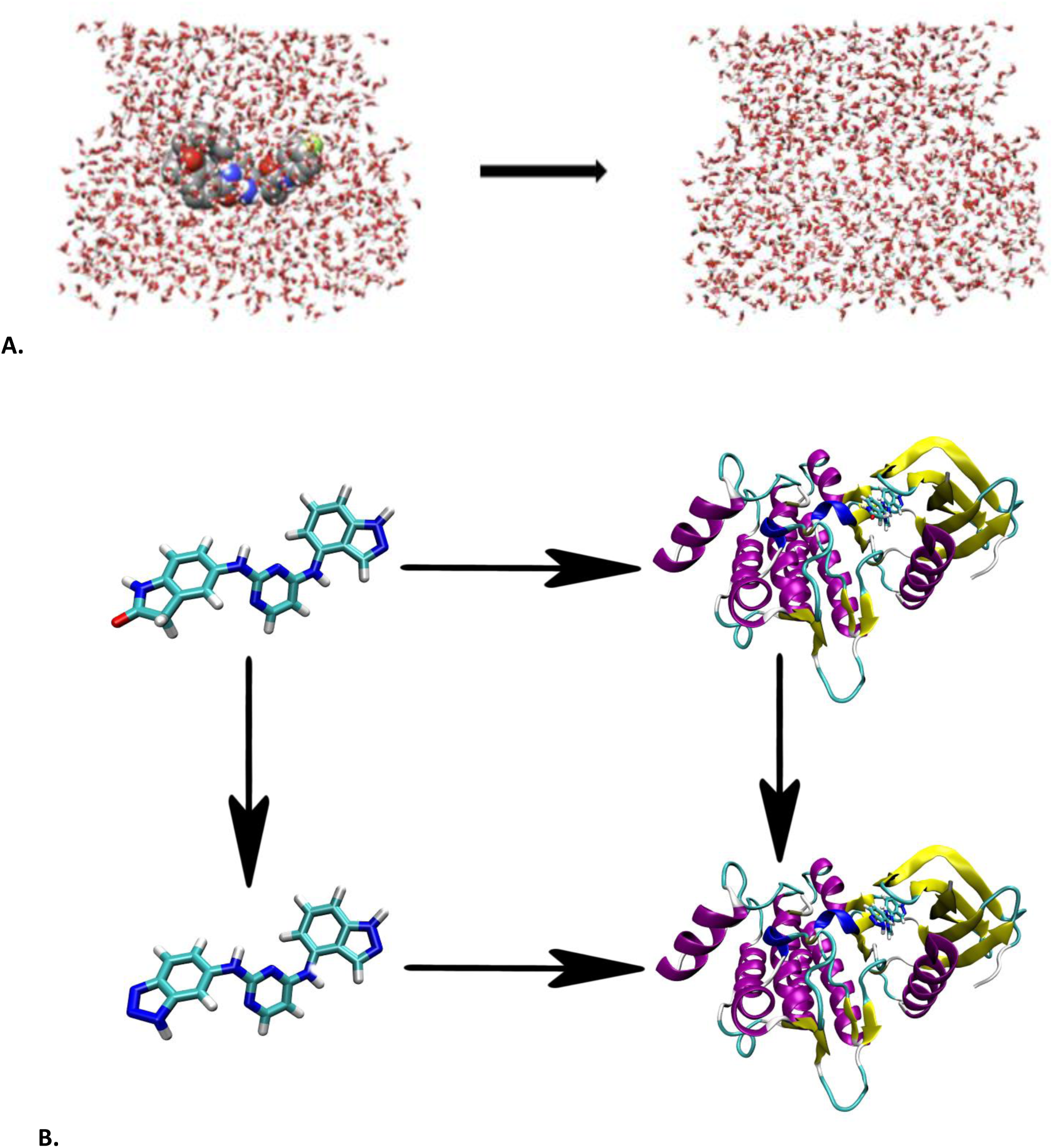
Perturbation cycle for sample calculations performed to test the accuracy and performance of our new GPU implementation of TI. Each molecule represents a thermodynamic endpoint of a calculation. **Top**: Solvation free energy of N1-((2S,3S,5R)-3-AMINO-6-(4-FLUOROPHENYLAMINO)- 5-METHYL-6-OXO-1-PHENYLHEXAN-2-YL)-N3,N3-DIPROPYLISOPHTHALAMIDE, a promising BACE-1 inhibitor.^14^ **Bottom**: Relative free energy of binding of ligands GTC000107A and GTC000112A to spleen tyrosine kinase (Syk) from the GSK Syk database available through the D3R project.^15^ The transformation includes 3 deletions and two atom changes from GTC000107A to GTC000112A.

### Solvation free energy

For our present purpose of demonstrating the precision of our GPU TI code compared to the existing CPU based TI implementation it was important to test calculations that were difficult to converge. In the original implementation of TI in pmemd,^16^ the solvation free energy calculation was the most difficult to converge, despite free energies of solvation (ΔG_solv_) having been one of the first types of free energy calculations attempted.^17^ In that instance, ΔG_solv_ was defined as decoupling the ligand from the water over the course of one simulation (**Figure 1A**). We chose a potential inhibitor of beta secretase-1 (BACE-1), N1-((2S,3S,5R)-3-AMINO-6-(4-FLUOROPHENYLAMINO)- 5-METHYL-6-OXO-1- PHENYLHEXAN-2-YL)-N3,N3- DIPROPYLISOPHTHALAMIDE as our test system here.^18^

### Relative free energy of binding

Relative free energy of binding (RFEB) calculations are the most widely used application of thermodynamic integration given their utility in early stage (lead discovery and lead optimization) small molecule drug discovery.^19,20^ For this very reason we have chosen to demonstrate an RFEB calculation in this paper. Our example thermodynamic cycle for RFEB is shown in figure 1B for the spleen tyrosine kinase (Syk) system. Syk was chosen because it has a stable, well-defined binding pocket with a crystal structure. Ligands GTC000107A and GTC000112A were chosen from the D3R database because they were shown to have similar binding poses^15^ These similar binding poses meant that convergence issues were less likely to dominate the results allowing us to better test the GPU implementation and to compare the differences in the precision model between the CPU and GPU implementations. As is typical in RFEB calculations, alchemical transformations from GTC000107A to GTC000112A were performed for the ligands bound to the protein and the ligands free in solution.

### Docking of ligands to Syk

Ligands were docked to crystal structure 1XBA^21^ using AutoDock Vina.^22^ Gaps in the crystal structure (residues 360-362, 393-394 and 405-406) were filled using Modeller 9v2.^23^ Side chains for residues 402, 420, and 448 were treated as flexible, while the rest of the protein was treated as rigid. Ligands were fully flexible.

## Simulation details

All the simulations follow a similar protocol; any details that are specific to a particular system will be described below

### System preparation and equilibration

The leap module in AMBER 16 was used to parametrize all systems. Protein models used the Amber ff14SB forcefield,^21^ with theTIP3P^22^ model for water. Neutralizing counterions, sodium or chloride, were added to each system as needed using TIP3P ions with parameters from Joung and Cheatham.^23,24^ Ligands were parametrized using the second generation generalized Amber forcefield (GAFF2)^25^ for the bonded and van der Waals parameters. Partial charges for ligands were obtained using RESP^26^ fitting for the electrostatic potentials, calculated using Gaussian^27^ at the Hartree – Fock/6-31G^⋆^ level of theory. A cubic periodic box was used with a minimum distance of 15 Å between any box edge and any solute atom. All systems were minimized for 1000 cycles of steepest descent followed by 1000 cycles of conjugate gradient. Solute atoms were restrained with a restraint weight of 10 kcal/(mol^⋆^Å)^2^. Minimization was followed, for all values of λ, by 100ps of heating at constant volume and then 1 ns of equilibration at constant pressure. Temperature was regulated via a Langevin thermostat set to a target temperature of 300 K and a collision frequency of 5.0 ps^-1^. Pressure was regulated using a Monte Carlo barostat with a target pressure of 1.0 atm and pressure relaxation time of 2.0 ps.

### Production

All production simulations were run in the NPT ensemble with a Langevin thermostat set to 300 K with collision frequency of 5.0 ps^-1^, and a Monte Carlo barostat at a target pressure of 1.0 atm and pressure relaxation time of 2.0 ps. The direct space cutoff was set to 10 Å for both van der Waals and electrostatics. Long range electrostatics we handled via the Particle Mesh Ewald (PME) method^28^ with a FFT grid spacing of ˜ 1 point per angstrom. Both the solvation free energy calculation and the RFEB calculation were run with 11 equally spaced lambda windows ranging from 0 to 1. The default values for scalpha (0.5) and scbeta (12.0) were used. Energies were printed every 0.5 ps. CPU simulations were run using pmemd.MPI from AMBER 16 with 12 Xeon Phi cores with a single socket running at 2.50 GHz on an Intel Haswell standard compute node on XSEDE Comet.^29^ Pmemd.MPI was compiled with MVAPICH2 2.1. GPU simulations were run on NVIDIA GeForce Titan-X GPUs with Pascal architecture using the current AMBER 18 development tree with our GPU TI support incorporated. The GPU code was compiled for the default SPFP precision model using CUDA 8.0 and NVIDIA driver version 367.57. Both pmemd.MPI and pmemd.cuda were compiled using the gnu compiler in gnutools 2.69. Simulations were run with a 1 fs time step (except for timing comparisons using H-mass repartitioning). The solvation free energy system was simulated for 15 ns. The RFEB complex and solvated systems were simulated for 10 ns. The first 5 ns of each simulation was discarded for equilibration purposes. The VDW change of the RFEB complex system required an additional 10 ns of simulation to converge. Each simulation was replicated three times with unique random seeds in each case.

### Single step alchemical change versus separate charge/VDW changes

In transformations involving charge changes, it is possible in AMBER to use a softcore coulomb potential alongside the softcore Lennard-jones potential. To test both approaches for reproducibility between the two codes, the solvation free energy calculation was run with softcore electrostatics, while the RFEB was run in separate steps, turning charges off linearly before performing VDW changes.

### H-Mass Repartitioning

Hydrogen masses bound to heavy atoms were repartitioned to 3.024 Daltons using ParmEd to allow for a 4-fs time step.^30^ Some short simulations were run with identical input parameters, except the additional use of SHAKE to restrain the bonds between hydrogen and heavy atoms. To obtain the same number of data points, print and write frequencies were quadrupled for these simulations.

### Performance comparison between CPU and GPU

All CPU simulations were run on using the pmemd.MPI module 12 Intel Xeon E5-2680v3 cores at 2.50 GHz on a single socket of an Intel Haswell standard compute node on XSEDE Comet. All GPU simulations were run on NVIDIA GeForce Titan-X (Pascal) GPUs with driver 367.57 and CUDA 8.0.

## Analysis

All integrations were carried out using a cubic spline over 11 lambda windows: 0.0, 0.1, 0.2, 0.3, 0.4, 0.5, 0.6, 0.7, 0.8, 0.9, 1.0. The Alchemical analysis python package^31^ was used to calculate our free energy estimates and associated errors. Autocorrelation times were estimated using pymbar^32^. Uncertainties for average values are the standard deviation of the replicate calculations.

## Results and Discussion

### Numerical comparison

Previous work by several groups has shown the CPU implementation of TI to be already capable of predicting experimental free energies.^33–36^ Thus for the purposes of validating our GPU implementation we consider values that agree within statistical error to the CPU implementation to indicate success.

The results of the solvation free energy and RFEB calculations are shown below (**Table 1**). The average results of the CPU and GPU simulations agree to within the standard deviation of three replicates.

**Table 1:** Numerical comparison of free energy of solvation and RFEB calculations on CPU and GPU. All values are the average ± the standard deviation of three replicates. RFEB – complex is the sum of the simulations in which the ligands are bound to the protein. RFEB – solution is the sum of the simulations in which the ligands are free in solution. Values for individual charge transformation and Lennard-Jones transformation simulation can be found in SI table 1.

| calculation | code | Relative Gibbs free energy change (kcal/mol) |
| --- | --- | --- |
| Solvation free energy | CPU | 53.0 ± 2.4 |
|  | GPU | 53.8 ± 2.4 |
| RFEB - complex | CPU | 27.50 ± 0.46 |
|  | GPU | 27.36 ± 0.55 |
| RFEB - solution | CPU | 28.95 ± 0.12 |
|  | GPU | 28.86 ± 0.05 |
| RFEB ($\Delta\Delta G$) | CPU | -1.45 ± 0.48 |
|  | GPU | -1.50 ± 0.55 |

The results agree between the CPU and GPU codes but, as shown with the performance numbers below, the GPU code obtained the result in approximately 1/30^th^ of the time it took to run the calculations on all CPUs cores within the node. For the RFEB calculation, the charge and VDW changes were carried out separately. These calculations also agreed to within statistical error (SI table 2).

Since the values measured here are ensemble averages using a Langevin thermostat that introduces random friction forces to control the temperature, it is still potentially possible that while the two codes agree within statistical error that we are using an incorrect potential in the GPU code due to some subtle implementation bug. It is thus useful, from a code validation perspective, to directly compare the potentials (both energy and gradients) between the CPU and GPU codes. **Table 2** below compares initial energies for the same starting structure between the CPU TI (DPDP precision model) and GPU TI (SPFP precision model).

**Table 2:** Numerical comparison of energies at step 0 of our RFEB trajectories. Results are for Lennard-Jones parameter change with softcore VDW potential. Electrostatic non-bonded includes both direct space and PME reciprocal space terms. “Bonded terms” includes bonds, angles, dihedrals, and 1-4 adjustment terms. DVDL, the derivative of potential with respect to lambda, is the sum of the contributions from all potential terms (bonded, VDW, and electrostatic). Forces were also equivalent to the same numerical precision, but are not shown due to the sheer number of atoms for which force is calculated.

| Term | CPU (kcal/mol) | GPU (kcal/mol) |
| --- | --- | --- |
| VDW non-bonded | 16209.8375 | 16209.8375 |
| Electrostatic non-bonded | -171344.5200 | -171344.5191 |
| Bonded terms | 19937.6161 | 19937.6162 |
| DVDL | -35.4512 | -35.4518 |

Energy and DVDL values agree to at least 6 significant figures which agrees with previous comparisons of
conventional MD simulations between the DPDP (CPU) and SPFP (GPU) precision models. This slight
difference is due to different charge representations between the CPU and GPU code. The CPU code
misrepresents the charges beyond the 6th decimal place on certain atoms.

## Performance comparison

The use of GPUs has allowed for a greater than 30X average performance increase over a single
socket 12 Intel Xeon E5-2680v3 core node. (Figure 2).

**Figure 2:**
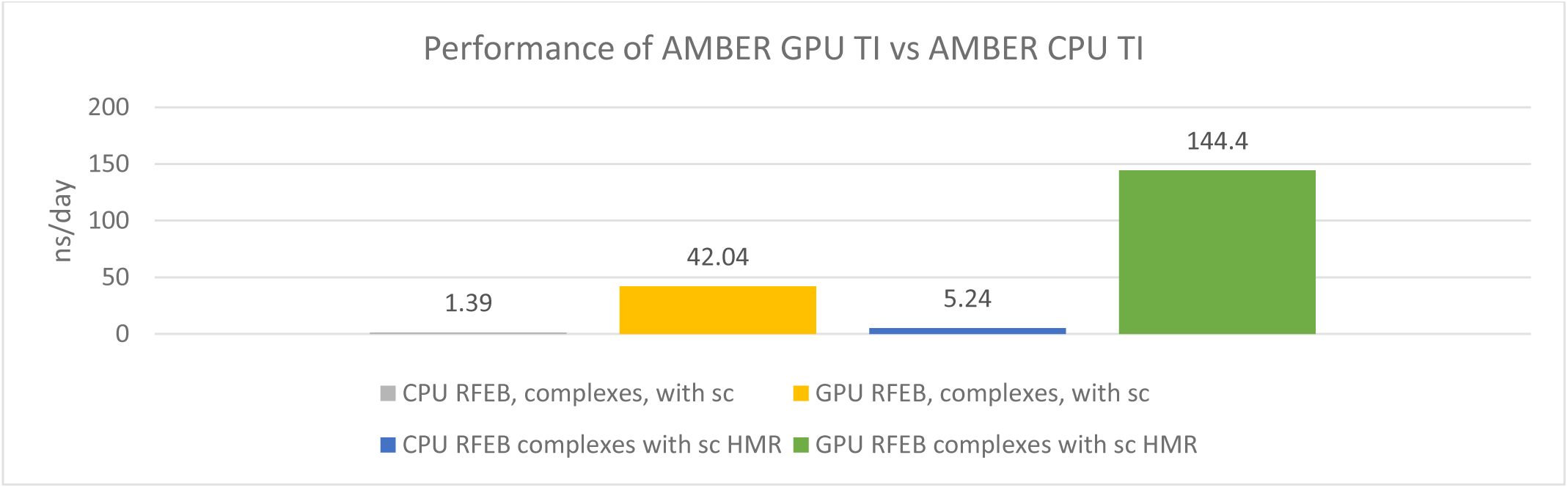
comparison of average performance of CPU and GPU TI code for a protein-ligand binding calculation. CPU code was run on 12 Intel Xeon E5-2680v3 cores, while the GPU code was run on a single NVIDIA GeForce Titan-X [Pascal] GPU. Numbers are from a VDW transformation from the lambda = 0 window. There was a total of 44,907 atoms, 81 of which were defined as TI atoms, with 3 being softcore. Charges on the softcore atoms were turned off. The GPU code performance did not scale as well with HMR compared with the CPU code (3.43X vs 3.77X) because of the higher energy printing frequency (every 125 steps) which more negatively affects GPU performance than CPU performance.

Hydrogen mass repartitioning enabled the use of a 4-fs time step. We noticed no issues in using 4-fs as a time step, in keeping with work done by others.^30,37^ Overall, we have achieved a greater than 360X performance enhancement for RFEB protein-ligand complex systems over a single CPU core, or greater than 30X performance improvement over a CPU node. With our code, these calculations can now be carried out on the order of hours on a single cost-efficient GeForce GPU, instead of weeks on a single node or having to utilize large number of CPU nodes with expensive interconnects. For way of reference the node used in this work can be purchased for approximately $2800 with a single GeForce Titan-X [Pascal] GPU. Our recommended configuration would, at the time of writing, be a system containing 2 x E5-2620V4 CPUs, 64GB of memory and 4 x NVIDIA GeForce 1080TI GPUs. This configuration currently costs between $5000 and $6500 and can run 4 lambda windows at once, one on each GPU, obtaining performance equivalent to that shown above for each window simultaneously.

## Conclusion

We have demonstrated GPU enabled TI in the AMBER molecular dynamics suite that will form the officially supported GPU TI implementation to be released in the upcoming AMBER v18. Our implementation compared with the most recent version of TI in pmemd8 is over 360X faster for protein-ligand binding than a single CPU core, 30X faster compared with a single node if using a single GPU per node and 120X faster per node if one considers that 4 GeForce GPUs can be added to a single node in a very cost effective manner. These performance improvements come without sacrificing precision or accuracy. The input format is identical to CPU pmemd, making the transition seamless for existing users. MBAR support has also been implemented and is in the process of being performance optimized. MBAR support will be available with the AMBER 18 release and the implementation will be discussed in follow up publications. The code described in this manuscript is available by request from the corresponding author as a patch against the release version of AMBER 17 and applicable updates as of Sept 1^st^, 2017.

## Acknowledgments

The authors thank Woody Sherman of Silicon Therapeutics, Thomas Fox of Boehringer-Ingelheim and Sasha Buzkho of Nant Biosciences for extensively beta testing the software and providing critical feedback. Daniel Mermelstein is supported in part by the Interfaces training program for multi scale biology from the National Institutes of Health (NIH). Work in the JAM group is supported in part by NIH, NBCR, and the NSF supercomputer centers. RCW’s contributions were funded by royalties received from licensing of the AMBER software by UCSF. No contributions to this work, monetary or otherwise, were made by NVIDIA Corp.

## Supporting info

**S1 Figure 1:**
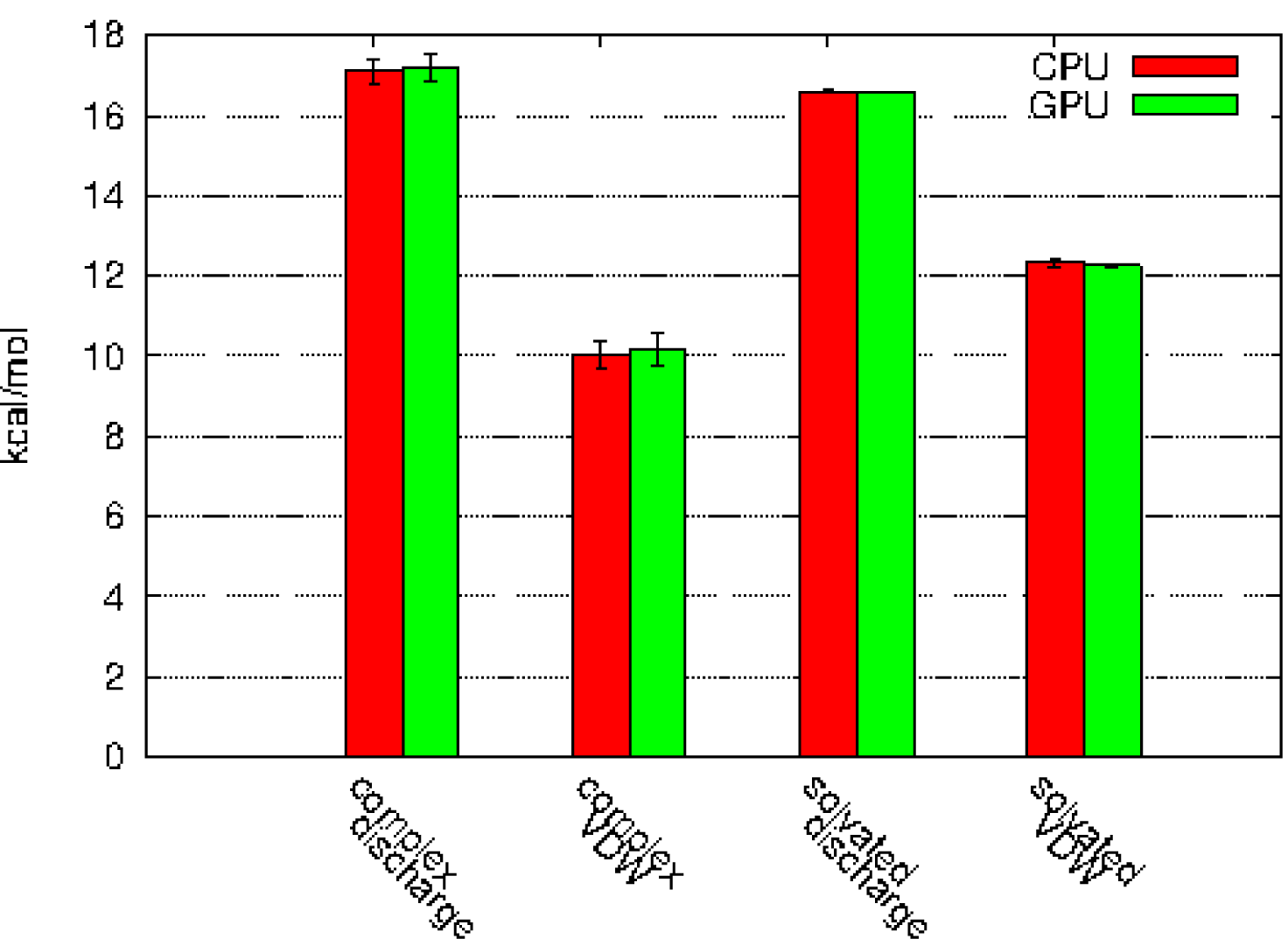
breakdown of RFEB calculation into charge change and Lennard Jones change. All caculated numbers agree within error (SI Table 1).

**S1 Table 1:** numerical values associated with SI figure 1.

| calc |  | step | Relative Gibbs free energy change (kcal/mol) |
| --- | --- | --- | --- |
| ligand binding - complex | cpu | discharge | 17.11 ± 0.29 |
|  | gpu | discharge | 17.21 ± 0.36 |
|  | cpu | vdw | 10.38 ± 0.36 |
|  | gpu | vdw | 10.15 ± 0.42 |
| ligand binding - ligands | cpu | discharge | 16.61 ± 0.01 |
|  | gpu | discharge | 16.59 ± 0.02 |
|  | cpu | vdw | 12.34 ± 0.11 |
|  | gpu | vdw | 12.27 ± 0.03 |

**S1 Figure 2:**
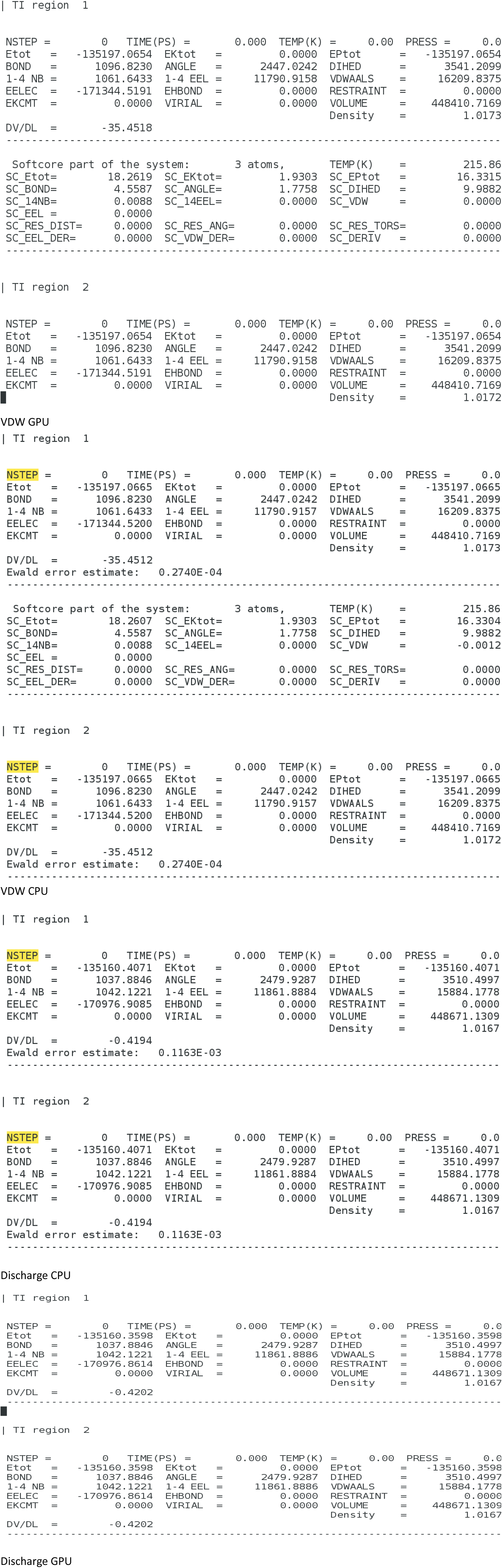
output initial energies for both Discharge and VDW changes in RFEB calculation.

